# Inducible activation of small GTPases reveals direct effector recruitment and signalling dynamics

**DOI:** 10.64898/2026.06.26.734842

**Authors:** Swati Singh, Marilyn Goudreault, Matthew J. Smith

## Abstract

RAS GTPases regulate cellular activity through the selective activation of effectors, yet identifying proteins directly recruited by small GTPases in cells remains challenging. Current approaches rely on extracellular stimulation or prolonged expression of constitutively active mutants, which trigger secondary signalling and negative feedback pathways. Most of the RAS superfamily also lack known upstream activators, limiting investigation of their biological functions. Here we develop SPaRTa (<u>S</u>equestered <u>P</u>rotein <u>a</u>ctivation through <u>R</u>AS-<u>T</u>EV <u>a</u>ctuation), an inducible system in which activated GTPases are maintained in a sequestered state by tethered effector-binding domains that can be released by rapamycin-induced reconstitution of split-TEV protease. We first applied SPaRTa to KRAS, as despite being one of the most intensely studied proteins in biology fundamental questions regarding its effector engagement remain unresolved. KRAS became activated within minutes of proteolytic release and stimulated rapid MAPK activity. Direct visualization of effector recruitment revealed distinct responses: BRAF was rapidly recruited to the plasma membrane (PM), whereas AFDN and RASSF5 accumulated gradually. In contrast, PI3Kα and SHOC2 were not recruited despite robust KRAS activation, though EGF stimulation efficiently re-localized PI3Kα to the membrane. This suggests that activation of KRAS alone is insufficient to stimulate PI3Kα. Sequestration shapes signalling as both AFDN and RASSF5 are subcellularly partitioned in the nucleus, and only prolonged KRAS activation re-located these effectors to the PM. Inducible activation of a designed RHOG SPaRTa resulted in ELMO1 recruitment and robust lamellipodia formation. Our system thus provides a broadly applicable platform for defining direct GTPase-effector interactions and signalling dynamics in cells.

## Introduction

RAS small GTPases function as molecular switches that regulate cellular physiology by cycling between inactive, GDP-bound and active, GTP-bound conformations. Activation promotes binding to downstream effector proteins, initiating signalling pathways that control proliferation, differentiation, adhesion, trafficking and more. Three classical RAS proteins (HRAS, KRAS and NRAS; hereon H/K/NRAS) are the best characterized members of the superfamily and signal through effectors including rapidly <u>a</u>ccelerated fibrosarcoma (RAF) kinases^1^, <u>p</u>hosphatidylinositol <u>3</u>-<u>k</u>inase (PI3K)^2^, <u>RAL</u> <u>g</u>uanine nucleotide <u>d</u>issociation <u>s</u>timulator (RALGDS)^3^, AFDN (Afadin/MLLT4/AF6)^4^, <u>R</u>AS <u>ass</u>ociation domain family <u>5</u> (RASSF5)^5^, <u>p</u>hospholipase <u>C e</u>psilon <u>1</u> (PLCε1)^6^ and the distinctive SHOC2 leucine rich repeat scaffold protein^7^. Through these effectors, H/K/NRAS convert extracellular signals into coordinated responses that direct cell proliferation. It follows that these GTPases are among the most frequently mutated proteins in human cancer^8^. Despite more than four decades of investigation, however, the mechanisms governing effector selection and signalling in cells remain incompletely understood.

Most effectors interact with activated H/K/NRAS through a conserved ubiquitin-like fold called a <u>R</u>AS-<u>b</u>inding <u>d</u>omain (RBD). Biochemical studies have revealed considerable differences in the affinity of individual RBDs, leading to the view that signalling is organized according to a hierarchy of effector binding^9^. BRAF and RAF1 bind RAS with nanomolar affinity^10,11^, whereas all other proposed effectors interact in the low µM range^12–17^. Signalling in cells, however, is unlikely to be governed by affinity alone. Effector recruitment will be influenced by protein abundance, compartmentalization, post-translational modifications (PTMs), competition, and the dynamic assembly of complexes on lipid membranes. Moreover, *in vitro* affinity measurements are typically performed using isolated GTPase and RBD domains, which excludes secondary contact sites in full-length effectors, PTMs, accessory proteins and lipid bilayers. Though biochemical studies have been invaluable for defining the molecular basis of GTPase-effector recognition, they cannot establish which effectors are directly recruited in cells. Thus, fundamental questions remain regarding which proteins are bound by activated GTPases, whether effectors exhibit distinct recruitment kinetics, and how signalling specificity is established.

The difficulty of studying GTPase signalling in cells extends beyond classical H/K/NRAS. The RAS superfamily comprises more than 160 GTPases^18^ that regulate processes as diverse as cytoskeletal remodelling, vesicular trafficking, polarity, and nuclear transport. While H/K/NRAS can be transiently activated through growth factor-mediated stimulation, most members of the superfamily lack defined physiological agonists or even known <u>g</u>uanine nucleotide <u>e</u>xchange factors (GEFs)^19^. Consequently, there are no approaches capable of acutely activating individual GTPases with sufficient temporal resolution to study early signalling events. A common solution has been to introduce mutations analogous to oncogenic KRAS variants and overexpress ‘constitutively active’ proteins for prolonged periods^20,21^. Such mutations have been biochemically validated for only a small subset of GTPases and, most importantly, prolonged overexpression will fundamentally rewire cellular signalling through sustained pathway activation and negative feedback^22–27^. Signalling and complex formation observed after hours/days of mutant expression is therefore unlikely to reflect direct and immediate consequences of GTPase activation.

Even for extensively studied H/K/NRAS, early signalling events following GTP loading remain difficult to investigate. Activation of H/K/NRAS using extracellular stimuli such as <u>e</u>pidermal <u>g</u>rowth factor (EGF) will also activate parallel pathways downstream of receptor tyrosine <u>k</u>inases (RTKs)^28^, making it challenging to distinguish proteins directly recruited by RAS from those requiring secondary signals. It thus remains unclear whether several proposed effectors, such as PI3K and SHOC2, are recruited directly by RAS, through RAS-independent signals, or by a combination. The duration of signalling also profoundly influences cellular behaviour. Transient or sustained activation of the MAPK pathway has long been recognized to produce distinct biological outcomes^29–32^, including proliferation versus differentiation, highlighting that signalling during the first minutes of H/K/NRAS activation may differ substantially from networks established after prolonged stimulation. An ideal experimental approach would therefore permit synchronized activation of an individual GTPase with minute-scale temporal resolution, allowing direct effector recruitment to be distinguished from secondary events.

To address these challenges, we developed SPaRTa (<u>S</u>equestered <u>P</u>rotein <u>a</u>ctivation through <u>R</u>AS-<u>T</u>EV <u>a</u>ctuation), an inducible platform that enables acute activation of RAS superfamily GTPases in living cells. Activated GTPases are maintained in an occluded, signalling-incompetent state through sequestration by a tethered RBD domain that is released by rapamycin-induced reconstitution of a split-TEV protease. Applying SPaRTa to KRAS enabled direct visualization of effector recruitment to the plasma membrane with temporal resolution. We could distinguish proteins recruited by activated KRAS alone from those requiring additional inputs and identified subcellular sequestration as a mechanism regulating effector availability. We show that inducible activation of RHOG leads to lamellipodia formation, illustrating SPaRTa is broadly applicable across the superfamily and provides a general strategy for investigating direct GTPase-effector interactions, signalling dynamics, and immediate cellular responses to GTPase activation.

## Results

### A split-TEV system for inducible and specific activation of small GTPases in cells

The study of direct signalling from activated RAS GTPases in cells is confounded by the requirement for prolonged expression of activated mutants. To circumvent this, we sought to design a system in which an activated GTPase is maintained in a sequestered state and released on a defined timescale. The core strategy tethers a low-affinity effector RBD domain to a mutationally activated GTPase using a flexible linker, physically occluding the G-protein from binding cellular effectors. The linker region encodes a protease recognition sequence that enables conditional, acute release upon cleavage. For this, we exploited a rapamycin (rap)-inducible split-TEV system in which the NIa protease from tobacco <u>e</u>tch <u>v</u>irus (TEV) is divided into two catalytically inactive fragments: nTEV (residues 1-118) fused to FRB and cTEV (residues 119-219) fused to FKBP^33^. The module reconstitutes active TEV protease upon rap-induced FRB:FKBP heterodimerization (**Fig. 1A**). TEV has no endogenous mammalian substrates, ensuring that all cleavage events are restricted to the engineered linker.

**Figure 1.**
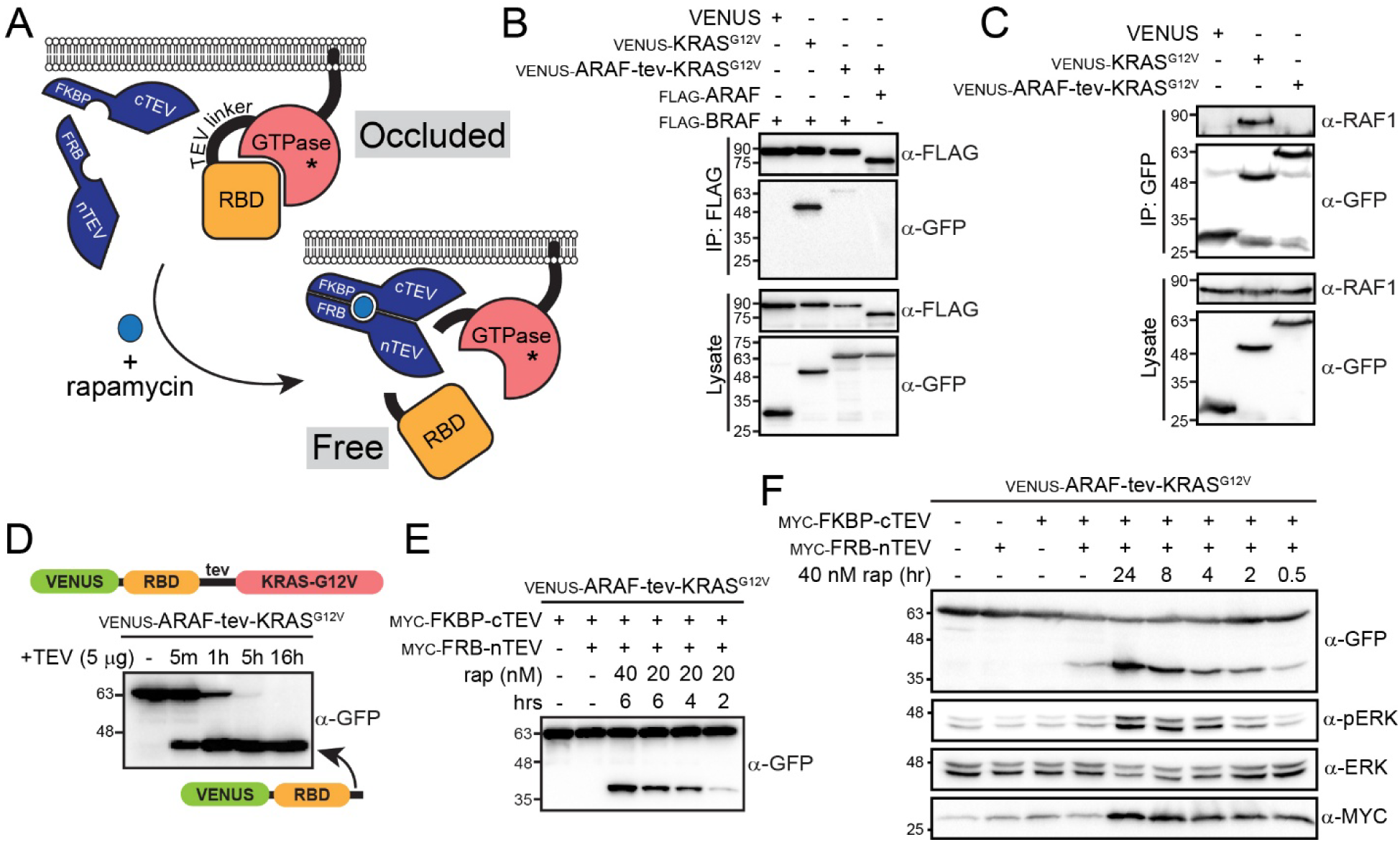
Development of a split-TEV approach to conditionally activate small G-proteins upon addition of rapamycin (rap) to cell culture media. (A) Schematic showing design of the inducible system. GTPases with activating mutations (*) are tethered to a moderate affinity effector binding domain (RBD) using a linker that includes a TEV protease recognition sequence. Occluded GTPases are co-expressed with FKBP-cTEV and FRB-nTEV, which reconstitute upon addition of rap to activate protease activity. Cleavage of the linker releases the RBD and free’s the activated GTPase for signalling to cellular effector proteins. (B) Tethering of RBD with the TEV-linker prevents binding of overexpressed ARAF and BRAF to activated KRAS^G12V^. Blotting for VENUS-tagged proteins following immunoprecipitation (IP) of FLAG-BRAF shows KRAS^G12V^ forms a complex, whereas a VENUS alone control or VENUS-ARAF-tev-KRAS^G12V^ do not co-precipitate with BRAF. The occluded GTPase is equally unable to complex with FLAG-ARAF. (C) Endogenous RAF1 does not co-precipitate with occluded KRAS^G12V^. A VENUS alone control, VENUS-KRAS^G12V^, or the designed VENUS-ARAF-tev-KRAS^G12V^ construct were precipitated and an anti-RAF1 blot used to resolve complex formation. Only KRAS^G12V^ binds RAF1, with neither VENUS alone nor the occluded GTPase associating with the endogenous kinase. (D) Release of the RBD from the occluded complex with exogenous, purified TEV. 5 µg of TEV protease was added directly to cell lysates expressing the intact, occluded GTPase protein and samples were incubated for the indicated timepoints. A lower MW band comprising VENUS-ARAF RBD is visible after 5 min of incubation with TEV. (E) Co-expression of occluded GTPase with cTEV and nTEV to assay rap-induced cleavage of the linker. Either 20 or 40 nM rap was added for the indicated timepoints (2-6 hours) to reconstitute TEV. A lower MW band comprising VENUS-ARAF RBD is observable even at 2 hours activation with 20 nM rap. (F) Full timecourse of occluded VENUS-ARAF-tev-KRAS^G12V^ activation induced by rap over 24 hours. The occluded GTPase was co-expressed with nTEV and cTEV and 40 nM rap added for the indicated timepoints. The protease fragments are tagged with MYC and are the same MW. Blotting for pERK/ERK reveals downstream signalling from KRAS^G12V^, only evident once the tethered RBD is released by TEV-linker cleavage.

To build and validate the first inducible GTPase we focused on KRAS, which has the most extensively characterized effector network with activating mutations and downstream readouts well established. We selected the RBD of ARAF as an occluding domain, as it binds KRAS with an affinity (dissociation constant (K_d_)) of approximately 1 µM^9,34^, sufficient for stable sequestration yet low enough that rapid dissociation is expected after proteolytic release. An N-terminal VENUS tag was incorporated to assist validation, and the archetype construct thus comprised VENUS-ARAF-tev-KRAS^G12V^ (the G12V mutation conferring constitutive activation). To establish whether the activated GTPase was occluded by the RBD, the construct was first co-expressed in HEK 293T cells with FLAG-tagged BRAF or ARAF, and complex formation assessed following anti-FLAG immunoprecipitation (IP). Free VENUS-KRAS^G12V^ alone was efficiently precipitated by FLAG-BRAF, whereas VENUS-ARAF-tev-KRAS^G12V^ did not associate with either BRAF or ARAF (**Fig. 1B**). This confirmed the tethered RBD effectively occludes the effector-binding site, preventing complex formation even with overexpressed partners. To establish that occlusion extends to endogenous effectors present at physiological concentrations, we performed a semi-endogenous co-IP from cells expressing VENUS alone, VENUS-KRAS^G12V^ or VENUS-ARAF-tev-KRAS^G12V^ and blotted for endogenous RAF1. The RAF1 kinase binds KRAS with an affinity in the low nanomolar range yet failed to co-precipitate with the occluded construct, while associating strongly with free KRAS^G12V^ (**Fig. 1C**). These data confirmed full occlusion of the GTP-bound KRAS protein, even against tight-binding effectors.

A prerequisite for this approach is that the TEV cleavage site embedded within the RBD:GTPase linker is accessible to protease. To test this, we first added purified TEV directly to cell lysates expressing the intact, occluded construct and monitored cleavage by the appearance of a low MW (molecular weight) VENUS-ARAF RBD product. Cleavage was detected within five minutes and was complete by 1-5 hours (**Fig. 1D**). This established the linker is fully solvent-accessible and that release of the RBD is rapid once protease is present. To test whether the inducible system in cells could cleave the linker, we co-expressed VENUS-ARAF-tev-KRAS^G12V^ with MYC-tagged FRB-nTEV and FKBP-cTEV. These split-TEV constructs were previously designed and used to selectively cleave cellular targets, particularly caspases^35,36^. We first treated cells with 20 or 40 nM rap over a timecourse spanning 2-6 hours. Cleavage was detectable at the earliest timepoint and increased progressively, with 40 nM rap producing more efficient cleavage (**Fig. 1E**). We extended the timecourse to cover 30 minutes-to-24 hours and stimulated with 40 nM rap (**Fig. 1F**). This showed a continuing accumulation of cleaved VENUS-ARAF RBD, with minimal non-specific cleavage in the absence of rap. These cells were starved overnight to permit detection of downstream signalling to the MAPK pathway. Indeed, we observed a progressive increase in pERK levels that paralleled formation of the cleavage product, demonstrating that release of KRAS^G12V^ from the tethered RBD is directly coupled to MAPK activation. Cells expressing the occluded construct without TEV fragments showed no pERK induction, confirming the signal is entirely attributable to released KRAS^G12V^. Systematic titration of rap from 10 to 100 nM confirmed all concentrations generate cleavage and pERK induction, but higher rap concentrations yield more efficient cleavage and increased pERK levels at earlier time points (**Fig. S1A**). Together, these experiments established a foundational occluded GTPase whereby mutationally activated KRAS^G12V^ is sequestered from its effectors by a tethered RBD and is released by rap-induced reconstitution of split-TEV protease, with downstream signalling directly proportional to the extent of proteolytic activation.

### Optimization of split-TEV kinetics and GTPase accessibility upon activation

The archetype system was functional but several aspects required optimization: 1) cleavage kinetics were too inefficient to resolve early effector recruitment events, as MAPK activation is typically evident 5 minutes after growth factor stimulation; 2) activating mutations are not equivalent and we sought to ensure all KRAS was GTP-bound following proteolytic release; 3) diffusion of the RBD upon cleavage is necessary to permit effector engagement and the affinity of the occluding domain must be considered. To increase the efficiency of TEV cleavage we reasoned that co-targeting reconstituted TEV to the PM, where KRAS constitutively resides *via* lipidation of its C-terminal CaaX motif, would increase local protease concentration at the substrate. To achieve this, we fused the 22-residue C-terminal membrane-targeting sequence of KRAS4B to the C-terminus of cTEV, generating cTEV-CaaX. Co-expression of FKBP-cTEV-CaaX with FRB-nTEV and occluded KRAS (VENUS-ARAF-tev-KRAS^G12V^) dramatically accelerated linker cleavage, producing nearly complete separation of the VENUS-ARAF RBD and KRAS^G12V^ modules and robust pERK within minutes of rap addition (**Fig. 2A**). Unfortunately, PM-targeted cTEV-CaaX also produced rap-independent cleavage, evident as accumulation of VENUS-RBD in untreated cells, indicating the TEV fragments were undergoing spontaneous reconstitution. Increasing or decreasing the rap concentration had no effect in this system (**Fig. S1B**). This confirmed that cTEV-CaaX, while greatly increasing cleavage kinetics, was not responsive enough for our inducible system.

**Figure 2.**
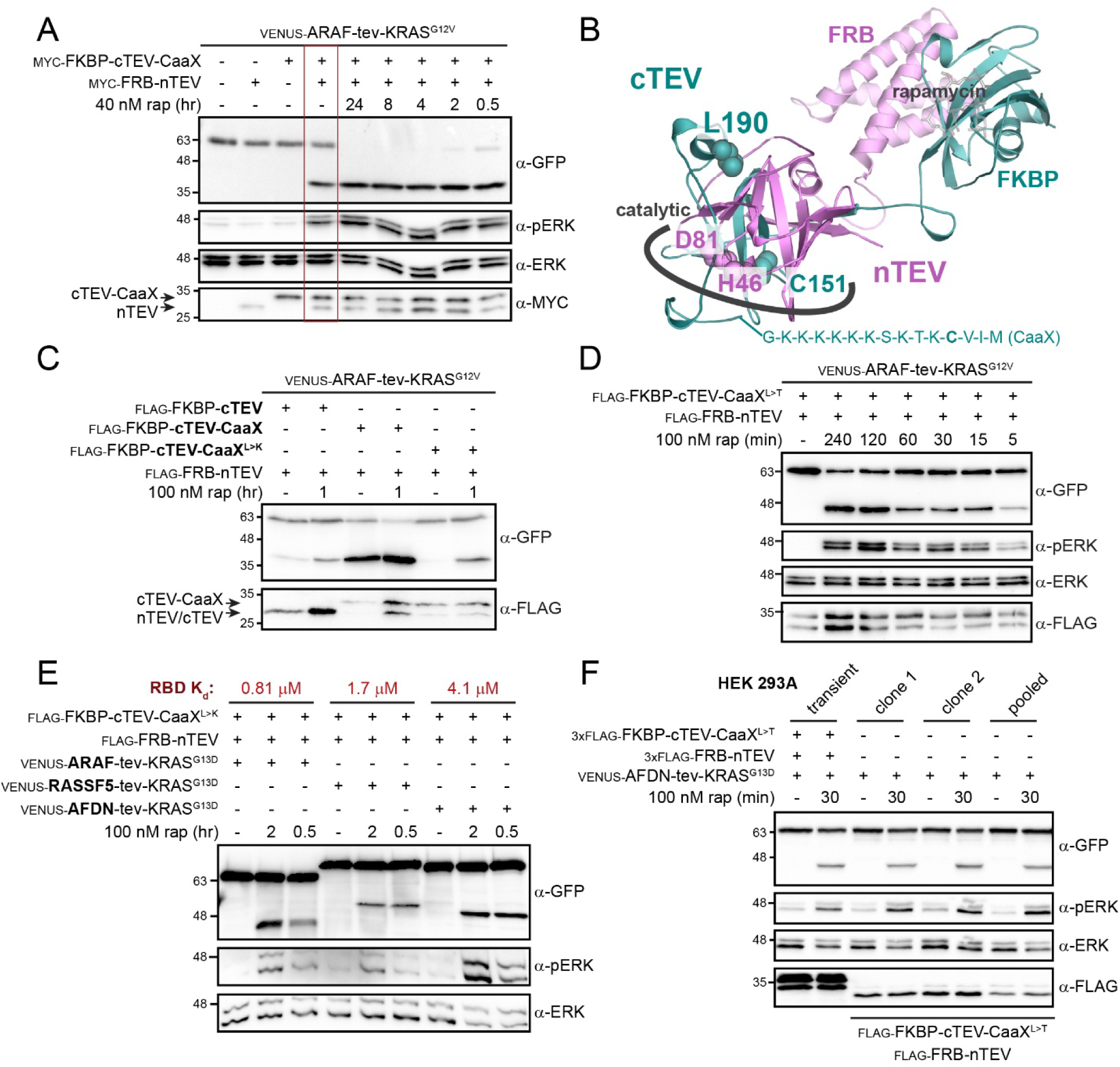
Optimization of the occluded GTPase assembly and TEV fragments. (A) Addition of a KRAS-derived CaaX sequence to the C-terminus of FKBP-cTEV. A timecourse of VENUS-ARAF-tev-KRAS^G12V^ cleavage over 24 hours following addition of 40 nM rap at the indicated timepoints was used to assay efficiency. Both VENUS-ARAF RBD (cleavage product) and pERK (downstream signalling) serve as reporters. An anti-MYC blot detects both cTEV and nTEV fragments (bottom). Cleavage using this system was observed even in the absence of rap (red box). (B) Structure model of the reconstituted FKBP-cTEV/FRB-nTEV protease with rap. The catalytic site comprises H46 and D81 from nTEV and the C151 residue of cTEV. At the reconstituted interface lies L190, the target of our mutational variants. The CaaX sequence from KRAS is added to the C-terminus of FKBP-cTEV. The model was constructed from existing structures PDBid 1Q31 (TEV) and PDBid 6M4U (FRB-FKBP with rapamycin). (C) Mutation at the TEV reconstitution interface restores sensitivity of membrane-targeted cTEV to rap. Either cTEV, cTEV-CaaX or cTEV-CaaX^L190K^ were co-expressed with nTEV and VENUS-ARAF-tev-KRAS^G12V^. Production of VENUS-ARAF RBD reports on cleavage efficiency in the presence or absence of rap. Expression of TEV fragments is verified in the anti-FLAG blot. (D) Cleavage timecourse of VENUS-ARAF-tev-KRAS^G12V^ by nTEV and the cTEV-CaaX^L190T^ variant spanning 5-240 min. pERK/ERK immunoblots report on KRAS activity, reflecting release of the occluding RBD domain by TEV. Expression of TEV fragments is confirmed by anti-FLAG blot. (E) Tethering of RBDs with diverse binding affinities for KRAS to optimize RBD diffusion following TEV-linker cleavage. ARAF (K_d_=0.81 µM), RASSF5 (1.7 µM) or AFDN (4.1 µM) RBDs were tethered to KRAS^G13D^ with the TEV-linker. Constructs were co-expressed with nTEV/cTEV-CaaX^L190K^ and 100 nM rap was added for the indicated timepoints. Presence of VENUS-tagged RBD proteins together with induction of pERK report on cleavage efficiency. The K_d_ of each interaction (measured previously by ITC) is shown at top in red. (F) Cell lines that constitutively express nTEV and cTEV-CaaX^L190T^ fragments. Both pooled (right lanes) and clonal (middle lanes) cell lines that stably express FLAG-TEV fragments were derived, and activity is compared with transient expression of 3x-FLAG nTEV/cTEV (left lanes). Cleavage of VENUS-AFDN-tev-KRAS^G13D^ is evaluated without rap or following addition of 100 nM rap for 30 min. VENUS-AFDN RBD and pERK report on efficiency.

To restore rap-dependence we considered mutations at the nTEV:cTEV-CaaX interface, informed by a prior computational study that identified cTEV L190 as a key determinant of reconstitution^37^. We moved both TEV fragments to expression vectors with N-terminal FLAG tags and generated cTEV-CaaX variants carrying L190K, L190T, L190A, L190I, or L190S substitutions. A structural model of reconstituted TEV shows L190 lies directly at the nTEV:cTEV interface and is distal to the catalytic triad residues H46, D81 (nTEV) and C151 (cTEV), rationalizing how a charge-introducing substitution at this position should modulate reassembly but not catalytic efficiency (**Fig. 2B**). A direct comparison of cTEV, cTEV-CaaX, and cTEV-CaaX^L190K^ confirmed that the L190K variant was entirely quiescent without rap and produced efficient rap-induced linker cleavage with no spontaneous reconstitution (**Fig. 2C**). This was also true of the L190T and L190A mutants, while the L190I substitution remained hyperactive and unresponsive to rap and the L190S mutant showed overall poor cleavage (**Fig. S2A**). A timecourse of occluded KRAS (VENUS-ARAF-tev-KRAS^G12V^) cleavage using nTEV and cTEV-CaaX^L190T^ revealed the PM-targeted variant is substantially more efficient, with both cleavage and pERK induction detectable within 5 minutes of rap stimulation (**Fig. 2D**). Thus, a membrane localization strategy combined with mutational inhibition of spontaneous reconstitution provided optimized split-TEV constructs for the inducible GTPase system.

Stimulation of the MAPK pathway, used to validate our approach, is generated from a high affinity interaction between KRAS-GTP and RAF kinases. We reasoned that binding to alternative effectors at weaker affinities could be more sensitive to the levels of released KRAS bound to GTP (i.e. in the activated conformation) and the capacity for effectors to compete with cleaved RBD. To ensure mutationally activated KRAS was fully GTP-loaded we considered the intrinsic nucleotide exchange rates of individual mutants. G12V KRAS exchanges GDP-to-GTP at a rate 1.8-fold slower than wild-type, while G13D KRAS confers rapid nucleotide exchange (15-fold faster than wild-type) that would ensure the GTPase remains fully GTP-loaded in cells^38^. We compared the two oncogenic mutants directly in the VENUS-ARAF-tev-KRAS background, and G13D regularly induced higher pERK levels than G12V (**Fig. S2B**) consistent with more sustained GTP occupancy. We assayed cTEV-CaaX^L190K^ cleavage and activation of an occluded KRAS^G13D^ construct against the archetype cTEV fragment (cytosolic), which further confirmed that PM-targeted TEV was more efficient at shorter timepoints (**Fig. S2C**). Notably, this experiment also compared acute activation of occluded VENUS-ARAF-tev-KRAS^G13D^ with prolonged expression of constitutively active KRAS^G12V^ (a standard approach to studying effector engagement in cells). The inducible system consistently produced higher pERK levels. This established that signalling from the occluded constructs is more analogous to growth factor stimulation and shows how prolonged mutant expression attenuates downstream signalling. To optimize the rate of GTPase release following proteolysis we compared three occluding RBD domains that bind KRAS with various affinities: ARAF (K_d_ = 0.81 µM)^9^, RASSF5 (K_d_ = 1.7 µM)^39^, and AFDN (K_d_ = 4.1 µM)^12^. Each was tethered to KRAS^G13D^, and cleavage was induced by addition of 100 nM (**Fig. 2E**) or 40 nM (**Fig. S2D**) rap. Efficiency was assessed by pERK induction and production of the RBD cleavage product. All constructs showed efficient linker cleavage, confirming that distinctive occluding domains do not impair TEV access, however, RASSF5 consistently attenuated pERK induction. This is likely due to an unusually slow RASSF5:RAS k_off_^13^, suggesting the RBD remains bound to KRAS for an extended period after cleavage. Both ARAF- and AFDN-occluded KRAS^G13D^ produced strong pERK responses within 30 minutes, with the AFDN variant performing at least as well despite its weaker K_d_. A full timecourse with VENUS-AFDN-tev-KRAS^G13D^ (**Fig. S2E**) showed this construct was efficiently cleaved and able to stimulate pERK production just minutes after addition of rap. The system was now optimized for rapid cleavage, GTP-loading, and diffusion of the occluding RBD.

To provide a stable experimental platform and avoid variability inherent to transient transfection, we derived monoclonal HEK 293A cell lines stably expressing FLAG-FRB-nTEV and FLAG-FKBP-cTEV-CaaX^L190T^. To confirm these cells were capable of cleavage and KRAS activation, we transfected VENUS-AFDN-tev-KRAS^G13D^ into a stable pool of cells expressing the TEV fragments, two clonal lines, and the parental HEK 293A line co-transfected with 3xFLAG nTEV and cTEV-CaaX^L190T^ (**Fig. 2F**). Following addition of 100 nM rap for 30 minutes, both VENUS-AFDN RBD production and pERK induction in the stable lines were comparable to the transient transfection, confirming the stable cells recapitulate inducible cleavage. We also assayed the full panel of occluding constructs, which confirmed the stable line replicates effector release rates observed in the transient system including the attenuated response of RASSF5-occluded KRAS (**Fig. S2F**). A truncated AFDN domain with only 17.8 µM affinity for RAS^12^ was also included, but showed MAPK activity in the absence of rap and was not carried forward. Thus, AFDN RBD occlusion, membrane-targeted cTEV-CaaX^L190T^, KRAS^G13D^, and stable TEV-expressing HEK 293A cells define an optimized system. This final configuration we refer to hereon as the KRAS SPaRTa (<u>S</u>equestered <u>P</u>rotein <u>a</u>ctivation through <u>R</u>AS-<u>T</u>EV <u>a</u>ctuation).

### Cell imaging reveals effector-specific recruitment dynamics to activated KRAS

An optimized SPaRTa should permit direct visualization of individual effectors recruited to the PM as an immediate and primary consequence of KRAS activation, free from secondary signals generated by growth factor:RTK engagement. To study this, we first focused on the high affinity interaction between KRAS and BRAF. We expressed CHERRY-BRAF in the HEK 293A stable line expressing nTEV/cTEV-Caax^L190T^, which revealed BRAF is predominantly cytosolic (**Fig. 3A**). In all experiments with KRAS, cells were serum starved overnight to prevent activation of endogenous RAS proteins. Interestingly, we did not observe a significant population of BRAF at the PM following 5-minutes of stimulation with EGF, consistent with past revelations that RAF membrane proximity is transient and comprises just a small fraction of cellular RAF^40^. We reasoned this was due to stoichiometry and repeated the experiment in cells with overexpressed, wild-type KRAS. In these cells, a 5-minute stimulation with EGF resulted in dramatic re-localization of CHERRY-BRAF to the PM (**Fig. 3A**), confirming our RAF construct could be efficiently PM-recruited by KRAS activation. We then assayed whether acute activation of KRAS SPaRTa could replicate growth factor-induced BRAF redistribution. Controls showed that co-expression of constitutively active and un-occluded VENUS-KRAS^G12V^ drives permanent BRAF localization at the PM, and that CHERRY-BRAF remains cytosolic upon expression of KRAS SPaRTa in the absence of rap (**Fig. 3B**). This confirmed the occluded GTPase is signalling-incompetent prior to proteolysis. Addition of rap for 30 minutes produced a complete and uniform redistribution of BRAF to the PM, and this localization was maintained through 4 hours of rap exposure. These results establish BRAF is a primary KRAS effector that is directly and rapidly recruited to GTP-loaded KRAS without requirement for any additional inputs.

**Figure 3.**
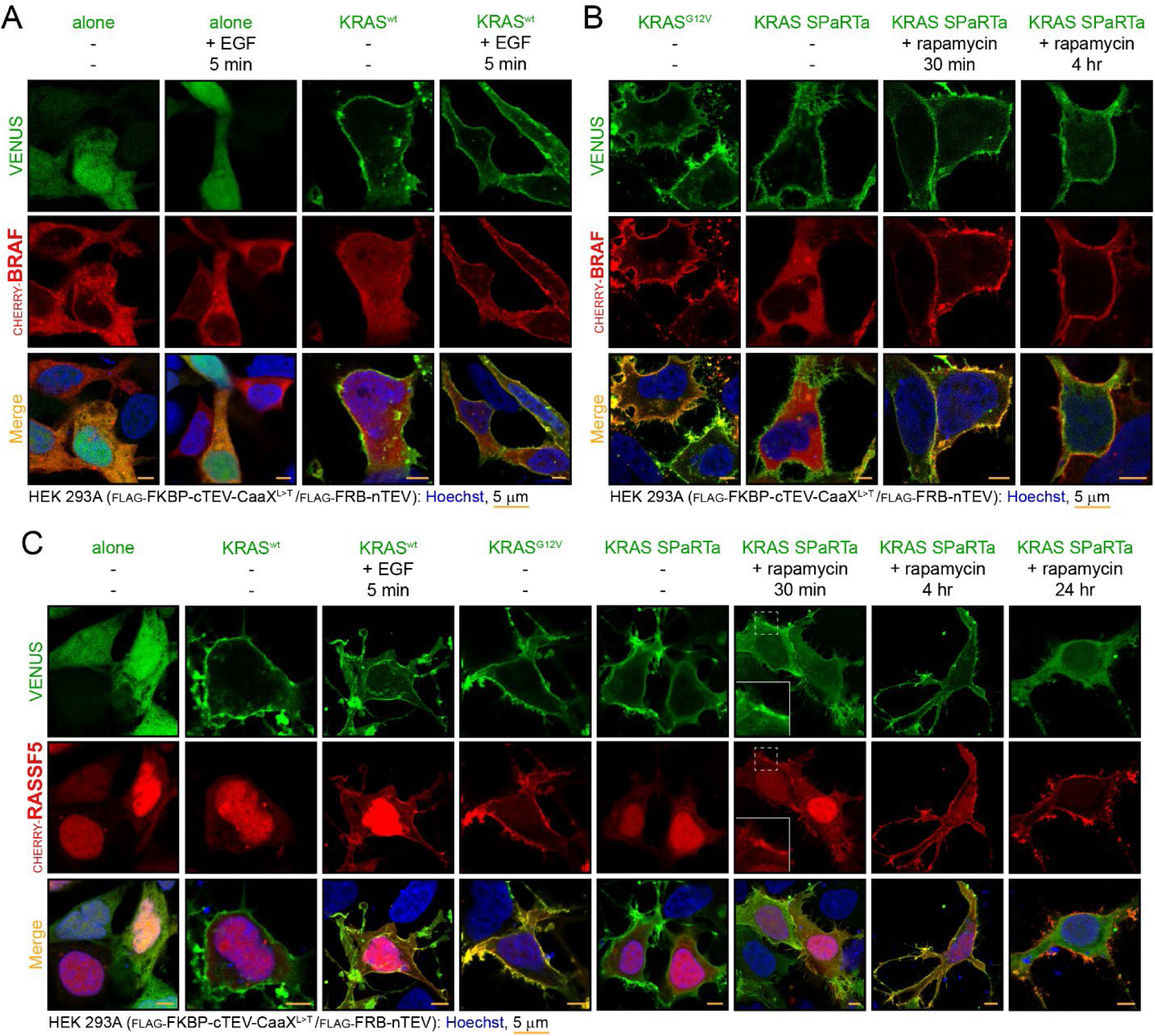
Recruitment of effectors to the PM by transient activation of KRAS SPaRTa. (A) Localization of CHERRY-tagged BRAF in starved HEK 293A cells. Confocal images show BRAF in the cytosol when co-expressed with VENUS alone or VENUS-KRAS^WT^. Addition of 50 ng/ml EGF for 5 min did not impact localization. CHERRY-BRAF is only observed at the PM of cells expressing VENUS-KRAS^WT^ stimulated with EGF for 5 min. Scale bar is 5 µm; Hoeschst-stained nuclei are blue. (B) Activation of KRAS results in redistribution of BRAF from the cytosol to the PM. Co-expression of CHERRY-BRAF with a constitutively activated mutant of KRAS (G12V) tagged with VENUS shows BRAF co-localized at the PM. Occluded VENUS-AFDN-tev-KRAS^G12V^ (SPaRTa) does not recruit BRAF in the absence of rap, but addition of rap for 30 min or 4 hr induces complete BRAF redistribution to the PM. Scale bar is 5 µm; Hoeschst-stained nuclei are blue. (C) CHERRY-RASSF5 gets recruited to the PM by EGF stimulation, constitutively active VENUS-KRAS^G12V^, or rap-induced activation of KRAS SPaRTa. Co-expression of RASSF5 with VENUS alone reveals its distribution in starved HEK 293A cells; predominantly nuclear. 50 ng/ml EGF stimulation for 5 minutes results in moderate RASSF5 distribution to the PM (when co-expressed with VENUS-KRAS^WT^), and 48-hour expression of VENUS-KRAS^G12V^ drives a complete PM recruitment of RASSF5. Activation of KRAS SPaRTa by addition of 40 mM rap for the indicated timepoints redistributes RASSF5 to the PM only with longer incubations (4-24 hrs). VENUS-AFDN RBD is observed in the cytosol following 24-hrs of cleavage. Scale bar is 5 µm; Hoeschst-stained nuclei are blue.

RAF kinases form high affinity complexes with KRAS but the capacity for activated H/K/NRAS to recruit alternative effectors in cells remains unclear. The KRAS SPaRTa system is ideal to resolve these questions, and we next studied how it impacts RASSF5, an effector that binds KRAS in the low µM range^39^. In starved cells, CHERRY-RASSF5 was predominantly nuclear with a fraction also evident in the cytosol (**Fig. 3C**), as has been observed previously^41^. Stimulation with EGF (in cells with overexpressed KRAS^WT^) produced only moderate RASSF5 redistribution to the PM, whereas 48 hours of VENUS-KRAS^G12V^ expression drove complete nuclear-to-PM re-localization. This suggested that sustained KRAS activity should be sufficient to overcome RASSF5 nuclear retention. Indeed, activation of the KRAS SPaRTa produced progressive RASSF5 redistribution to the PM, but its complete re-localization required extending rap stimulation to 4-24 hours. Released VENUS-RBD was clearly evident in the cytosol at the 24-hour timepoint, confirming ongoing TEV protease activity. To verify that effector recruitment was dependent on SPaRTa we used CHERRY-RASSF3 as a negative control, this effector being highly similar to RASSF5 but unable to directly bind KRAS^39,41^. RASSF3 remained entirely cytosolic and unresponsive under all conditions at every timepoint examined (**Fig. S3A**). These data demonstrate that RASSF5 is a direct and specific KRAS effector whose PM recruitment is kinetically limited by nuclear sequestration, requiring sustained GTPase activation to accumulate at the membrane.

Numerous additional candidate RAS effectors and regulatory proteins could be assayed in this system, and we chose three for which direct RAS binding and recruitment requires improved validation. Like RASSF5, AFDN partitioned between the nuclear and cytoplasmic compartments in starved cells (**Fig. 4A**). We applied the same series of activation conditions in cells expressing CHERRY-AFDN, and both EGF stimulation and co-expression with constitutively active KRAS^G12V^ redistributed the effector to the PM. rap-induced SPaRTa activation was also able to recruit AFDN, confirming direct KRAS activation alone is sufficient to alter AFDN subcellular distribution. AFDN responded to SPaRTa activation at prolonged timepoints, consistent with a less accessible cytoplasmic pool and requirement for nuclear exit. Whether PI3Kα and SHOC2 are direct KRAS effectors in cells is an important open question, and we could distinguish if KRAS activation alone or additional membrane-proximal inputs stimulated by growth factors are required. We monitored CHERRY-tagged PI3Kα and SHOC2 localization following either EGF stimulation, prolonged co-expression with activated KRAS^G12V^, or SPaRTa activation. Neither constitutively active KRAS^G12V^ nor SPaRTa release at any timepoint up to 4 hours produced a detectable change in PI3Kα (**Fig. 4B**) or SHOC2 (**Fig. S3B**) localization, with both remaining cytosolic under all conditions. Interestingly, EGF stimulation of cells co-expressing CHERRY-PI3Kα and VENUS-KRAS^WT^ produced rapid and robust PI3Kα recruitment to the PM within 5 minutes, confirming the effector is fully competent to re-localize and that failure to respond to KRAS activation alone reflects a genuine mechanistic requirement. SHOC2 remained cytosolic even upon EGF stimulation, indicating its association with RAS signalling components does not involve direct PM recruitment under any of our examined conditions. These data establish that neither PI3Kα nor SHOC2 membrane recruitment is driven solely by GTP-loading of KRAS, and that PI3Kα engagement at the PM requires additional signals generated by growth factor stimulation.

**Figure 4.**
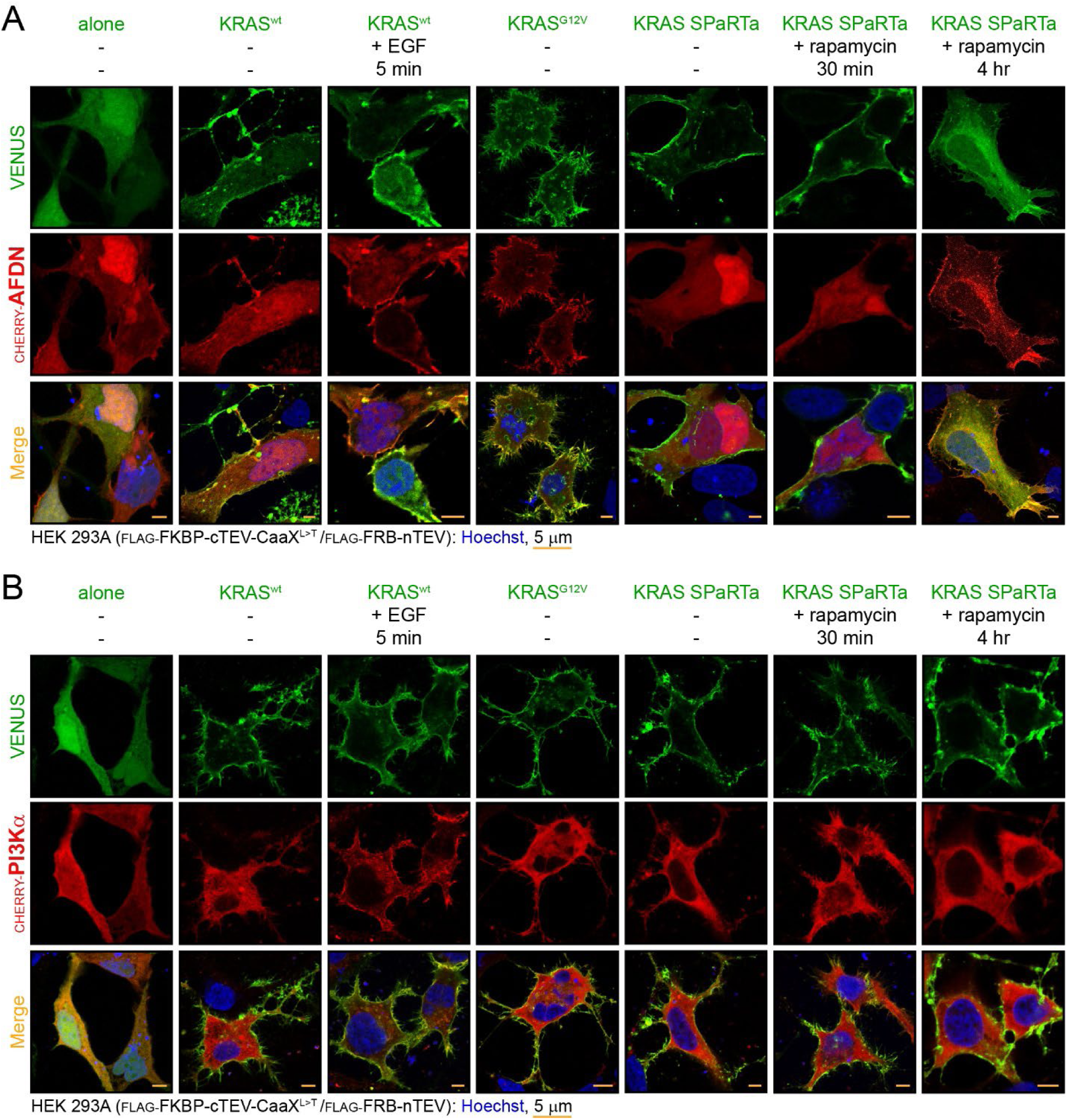
AFDN is recruited to the PM by KRAS SPaRTA, but PI3Kα remains in the cytosol. (A) CHERRY-AFDN has a nucleo-cytoplasmic localization in starved HEK 293A cells expressing a VENUS alone control. AFDN is recruited to the PM by EGF stimulation (50 ng/ml for 5 minutes in cells expressing VENUS-KRAS^WT^), constitutively active VENUS-KRAS^G12V^, or rap-induced activation of KRAS SPaRTa. Activation of SPaRTa was done by addition of 40 mM rap for the indicated timepoints. Scale bar is 5 µm; Hoeschst-stained nuclei are blue. (B) Only the addition of growth factor (stimulation with 50 ng/ml of EGF for 5 minutes in cells expressing VENUS-KRAS^WT^) redistributes CHERRY-PI3Kα from the cytosol to the PM. Neither co-expression of PI3Kα with constitutively active VENUS-KRAS^G12V^ nor activation of KRAS SPaRTa for the indicated timepoints (40 mM rap for up to 4 hrs) had any impact on PI3Kα localization. Scale bar is 5 µm; Hoeschst-stained nuclei are blue.

Quantifying the proportion of effector populations recruited to the PM by microscopy is challenging, as conventional co-localization metrics such as Pearson coefficients perform poorly when one binding partner is membrane-localized^42,43^. This problem is compounded in our SPaRTa system as cleaved VENUS-RBD domains transit into the cytosol following cleavage. We therefore attempted subcellular fractionation as an orthogonal biochemical approach to measure steady-state membrane association. For both BRAF and RASSF5, enrichment in the membrane fraction was detected exclusively in cells overexpressing KRAS^G12V^ for 48 hours; neither transient EGF stimulation nor SPaRTa activation at any timepoint produced measurable membrane enrichment of these effectors despite confocal imaging showing PM localization under the same conditions (**Fig. S4A/B**). We even attempted chemical crosslinking with DSS (disuccinimidyl suberate) prior to fractionation, intended to stabilize transient membrane interactions before lysis. This also failed to shift RASSF5 into the membrane fraction (**Fig. S4C**), though the crosslinker did generate significant laddering indicative of higher order complexes. Taken together, the fractionation approach supports previous suggestions^40^ that effector recruitment to the PM following acute KRAS activation is transient and dynamic, and this is readily resolved by confocal imaging but occurs at a frequency too low to be captured by bulk biochemical fractionation. This further substantiates the SPaRTa system as being an acute-phase GTPase activator, more analogous to growth factor-induced activation than prolonged overexpression of constitutive mutants.

### Extending the SPaRTa system to activation of the small GTPase RHOG

While classical H/K/NRAS GTPases can be activated by the addition of growth factors to media, most RAS superfamily GTPases lack defined agonists and cannot be acutely activated by extracellular stimuli. This severely limits investigation of immediate downstream interactions, and we sought to determine whether SPaRTa could be adapted for a non-RAS subfamily member. RHOG is a small GTPase in the RHO subfamily that activates ELMO/DOCK proteins to stimulate GDP-to-GTP exchange of RAC1 and subsequent cytoskeletal remodelling^44^. An RBD-like domain from the N-terminus of murine Elmo2 binds RHOG with a Kd of 7.8 µM^45^, which provided an optimal occluding domain. We generated a VENUS-Elmo2-tev-RHOG^G12V^ SPaRTa construct and verified sequestration by co-IP with CHERRY-ELMO1. ELMO1 did not co-precipitate with wild-type RHOG, VENUS alone, or the RHOG SPaRTa in the absence of rap, but formed a robust complex with constitutively active VENUS-RHOG^G12V^ (**Fig. 5A**). This confirmed that the Elmo2 RBD effectively occludes the RHOG effector-binding site prior to cleavage. To confirm that split-TEV activation kinetics are preserved, we assessed rap-induced cleavage of RHOG SPaRTa in the stable HEK 293A line (FLAG-FRB-nTEV and FLAG-FKBP-cTEV-CaaX^L190T^) alongside the KRAS SPaRTa as control. Addition of rap at 40 nM produced time-dependent accumulation of the VENUS-Elmo2 RBD cleavage product with an efficiency comparable to that of KRAS SPaRTa (**Fig. 5B**), confirming cleavage kinetics are governed by the TEV reconstitution mechanism and are indifferent to the identity of the occluded GTPase. Thus, the system is amenable to activation of alternative RAS superfamily GTPases, provided an occluding domain with affinity in the low micromolar range is available for sequestration.

**Figure 5.**
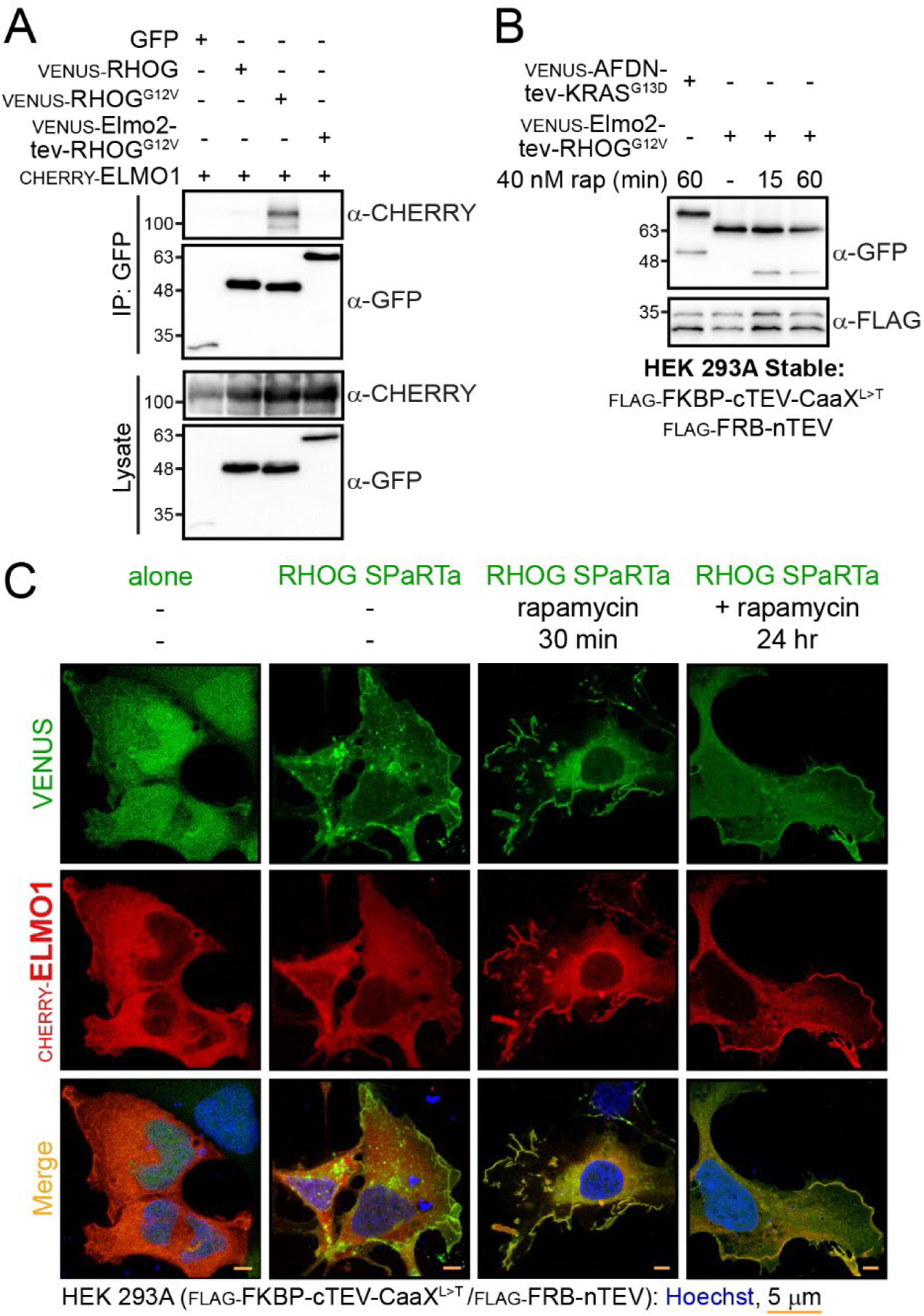
Assembly of a RHOG SPaRTa able to generate lamellipodia upon activation. (A) A VENUS-Elmo2-tev-RHOG^G12V^ construct effectively occludes the activated GTPase from binding its effector, ELMO1. HEK 293A cells co-expressing CHERRY-ELMO1 and either VENUS alone, VENUS-RHOG^WT^, VENUS-RHOG^G12V^ or the RHOG SPaRTa (in the absence of rap) were lysed and immunoprecipitated with anti-GFP. Only constitutively active RHOG^G12V^ co-precipitates with ELMO1. (B) Timecourse of occluded VENUS-Elmo2-tev-RHOG^G12V^ activation induced by rap addition over 60 min. Occluded RHOG was expressed in the HEK 293A stable line (nTEV and cTEV-CaaX^L190T^) and 40 nM rap added for the indicated timepoints. Cleavage of the KRAS SPaRTa (VENUS-AFDN-tev-KRAS^G13D^) was included as a control. Protease fragments are tagged with FLAG and expression is confirmed by an anti-FLAG blot. (C) Activation of RHOG SPaRTa results in redistribution of CHERRY-ELMO1 from the cytosol to the PM. Expression of VENUS alone with ELMO1 reveals its general localization in the cytosol Occluded VENUS-Elmo2-tev-RHOG^G12V^ does not recruit ELMO1 to the PM in the absence of rap, but addition of rap for 30 min induces its redistribution. Extensive lamellipodia are observed following 24-hrs of rap stimulation. Scale bar is 5 µm; Hoeschst-stained nuclei are blue.

To determine whether acute RHOG activation is sufficient to recruit ELMO1 to the PM and drive downstream cytoskeletal changes, we imaged cells co-expressing CHERRY-ELMO1 and the RHOG SPaRTa before and after stimulation with rap. ELMO1 was distributed diffusely throughout the cytosol in the absence of rap, with no evidence of PM localization (**Fig. 5C**). Addition of 40 nM rap for 30 minutes produced a clear redistribution of ELMO1 from the cytosol to the PM, suggesting an activated RHOG^G12V^:ELMO1 complex was being formed at the membrane. Extending the cleavage over 24 hours drove extensive lamellipodia formation consistent with downstream RAC1 activation^46^ (**Fig. 5C**). SPaRTa proteolytic cleavage over the timecourse could even be verified using the small number of cells remaining in tissue culture wells after removal of the cover slip (**Fig. S5A**). These results establish that SPaRTa generalizes to RHO subfamily GTPases with no requirement for modifying the core split-TEV machinery, providing a broadly applicable strategy for investigating GTPase-effector interactions and immediate signalling outputs across RAS superfamily members for which no acute activation approach exists.

## Discussion

SPaRTa addresses a fundamental limitation in the study of RAS GTPase signalling: the inability to acutely activate a singular GTPase in cells without engaging additional signalling machinery (e.g. RTKs stimulated by mitogens) or imposing adaptive changes that accumulate during prolonged expression of constitutively GTP-bound mutants. By sequestering a mutationally activated GTPase behind a tethered RBD and releasing it through rap-induced reconstitution of TEV, SPaRTa enables GTPase activation on timescales relevant to endogenous signalling. In its current configuration only a fraction of the total expressed SPaRTa constructs become cleaved. We were ultimately satisfied with this, as it more closely mimics endogenous RAS behaviour than the saturating activation produced by constitutive expression. The balance between cleavage efficiency and suppression of spontaneous reconstitution could be further refined, however, through additional interface mutagenesis or linker optimization. The platform is already well-suited to address numerous research questions, including dissecting effector-specific outputs from distinct oncogenic KRAS mutants, studying GTPases lacking known physiological activators, comparing effector recruitment kinetics, and identifying effectors whose engagement requires signals beyond activation of individual RAS proteins alone.

Constructing a SPaRTa for additional GTPases requires only 1) a constitutively activating mutation and 2) an occluding RBD binding in the 1–10 µM affinity range. This binding is tight enough for sequestration but weak enough for rapid post-cleavage diffusion. The attenuated pERK response of RASSF5-occluded KRAS despite efficient linker cleavage illustrates the consequence of exceeding this window, as the slow k_off_ of the RASSF5:RAS interaction^13^ appeared to delay effector access after proteolysis. The cTEV-CaaX targeting sequence can be substituted with any characterized membrane-targeting motif to redirect split-TEV reconstitution to the compartment where a GTPase resides, extending SPaRTa to those at endomembranes, Golgi, mitochondria, or endosomes without modification to any other component. Two limitations are worth noting: the system is irreversible, precluding experiments requiring GTPase inactivation, and the multi-domain architecture leaves limited space for additional modules (such as a TurboID tag for proximity-based proteomics), though this is likely feasible with careful engineering.

A key biological finding from applying SPaRTa to KRAS is that PI3K recruitment to the PM is not driven by GTP-loaded KRAS alone, instead requiring receptor-proximal signals generated by growth factor stimulation. The failure of PI3Kα to respond to either constitutively active KRAS^G12V^ or acute SPaRTa activation, contrasted with its rapid PM-recruitment following EGF stimulation, directly resolves an ambiguity that could not be addressed using mitogenic stimulation alone. The situation with SHOC2 is more complex; its nucleating GTPase remains debated^47^ with evidence implicating both MRAS and KRAS^48,49^. Our data show SHOC2 is unresponsive to growth factors or KRAS activation at the PM, consistent with earlier reports that wild-type SHOC2 does not translocate to the PM upon EGF stimulation^50^. Others have reported that upon EGFR activation, endogenous SHOC2 translocates to a subset of late endosomes containing RAB7^51^, but this was not resolved in our work. Ultimately, whether SHOC2 recruitment is nucleated by MRAS, KRAS, or another small GTPase remains an open question that SPaRTa is well positioned to address. Imaging experiments revealed that nuclear sequestration of RASSF5 and AFDN imposes a kinetic threshold on their PM recruitment that is determined by availability in the cytosol. This separation suggests nuclear retention functions as a rheostat: signals that promote nuclear export of either protein would expand the cytoplasmic pool and lower the threshold for KRAS-driven membrane recruitment, whereas signals reinforcing nuclear retention would dampen effector engagement even when the GTPase is fully active. The identities of the signals that control this nuclear-cytoplasmic partitioning, and whether they are coordinated with upstream RAS activation, represent important open questions.

The transient and dynamic nature of effector recruitment to active KRAS at the PM, demonstrated here by the discordance between confocal imaging and subcellular fractionation, is supported by a detailed study of endogenous fluorescently tagged RAF1 in EGF-stimulated cells^40^. In that work, only ∼10% of total RAF1 translocated to the PM following EGF stimulation, and following an early burst of translocation membrane-associated RAF1 was undetectable by both microscopy and subcellular fractionation, estimated at fewer than 200 molecules per cell. The failure of fractionation to capture this pool, despite clear functional evidence of RAF1 engagement with RAS-GTP at the membrane, precisely mirrors our observations with BRAF and RASSF5 in the SPaRTa system and reinforces the conclusion that bulk biochemical membrane enrichment assays are fundamentally unsuited to detecting the transient, low-stoichiometry interactions that characterize physiological effector recruitment.

The most consequential future applications of SPaRTa are likely to involve GTPases for which no physiological activator has been identified, a category that encompasses a large fraction of the ∼160 RAS superfamily members including ARF, RAB, RAS, and RHO proteins whose upstream regulation is poorly characterized. We have demonstrated the feasibility of this using RHOG, a small GTPase for which the activating G12V mutation has been characterized and an occluding RBD in the correct affinity range^45^ was available. Activation of the RHOG SPaRTa resulted in ELMO1 membrane recruitment and lamellipodia formation, and the system is primed to study kinetics of downstream RAC1 activation, the identity of DOCK proteins complexed with ELMO, and more. Our approach can also be used to address ambiguities in signalling research, particularly when *in vitro* biochemical results do not correlate with those from cells. This is true of GTPases from the RAP subfamily, which bind RAF RBD domains *in vitro*^52^ but do not activate the MAPK pathway to the same level as classical RAS proteins^53^. We used the dual-specificity AFDN RBD^54^ to generate a RAP1 SPaRTA, and indeed observed little pERK induction upon rap-stimulated cleavage (**Fig. S5B**). This further illustrates the importance of choosing downstream readouts when assaying the successful activation of a given SPaRTa/GTPase. For classical H/K/NRAS, an important near-term application will be direct comparison of oncogenic mutants (e.g. G12C, G12D, G12V, G13D, Q61L, etc.) for their capacity to engage effectors with unique kinetics or stoichiometries, a question confounded by prolonged expression as individual mutants could activate negative feedback at different rates. SPaRTa equalizes the activation time course and isolates intrinsic differences in effector engagement from secondary adaptive differences. Conversely, it will be important to compare the recruitment kinetics of distinct effectors to the same activated GTPase simultaneously, which would allow a hierarchy of effector engagement to be determined in real time.

We provide here a general strategy for temporally controlling the activation of individual RAS GTPases, uncoupling GTP-loading from the complex upstream signalling events that typically accompany activation. Beyond defining direct effector interactions, the SPaRTa platform should ultimately allow time-resolved proteomics, comparison of disease-associated GTPase mutants, characterization of poorly understood RAS superfamily GTPases, and evaluation of pharmacological signalling modulators. By providing temporal control over individual GTPases in cells and decoupling GTP-loading from upstream signalling, SPaRTa enables the earliest molecular events following small GTPase activation to be observed directly for the first time.

## Materials and Methods

### Plasmid constructs and antibodies

To generate the initial KRAS SPaRTa, the full sequence of human *KRAS4B* (GeneID 3845) was cloned into Gateway Entry vector pENTRA1A. Sequence encoding the RBD domain from ARAF (aa 18-93; GeneID 369) was inserted upstream, and a primer-encoded linker of 36 residues containing a TEV recognition site (ENLYFQS) was introduced between the ARAF and KRAS4B coding sequences. The consequent ARAF-tev-KRAS Entry clone was recombined into a mammalian expression Gateway Destination vector with an N-terminal VENUS tag using LR clonase (Invitrogen). To generate constructs with variable RBD affinities, the ARAF sequence was replaced with either RASSF5 (aa 205-363; GeneID 83593) or AFDN (aa 3-136; GeneID 4301). A shorter AFDN RBD with further reduced affinity^12^ was also used (aa 36-136). To generate the RAP1A SPaRTa, the KRAS4B coding sequence in the AFDN-occluded construct was replaced with that of *RAP1A* (GeneID 5906). For inducible activation of RHOG, the RBD of murine Elmo2 (aa 1-80; GeneID 140579) was cloned upstream of the TEV linker followed by the coding sequence for human RHOG (GeneID 391). For reconstitution of TEV protease, the original MYC-tagged FRB-nTEV and FKBP-cTEV were a kind gift from Dr. Jim Wells (UCSF; Addgene #100095 and #100096)^35,36^. These were PCR amplified, cloned into Gateway Entry vectors, and subsequently shuttled with LR clonase into destination vectors for either transient or lentiviral-based expression of the proteins in cultured cells with N-terminal FLAG or 3xFLAG tags. To add a membrane-targeting CaaX motif, sequence encoding the 22 C-terminal residues of human KRAS4B was inserted downstream of FKBP-cTEV. The panel of effectors as well as plasmids for expressing standalone GTPases were made by Gateway recombination of Entry clones into mammalian expression vectors with N-terminal CHERRY or FLAG tags. This includes clones for human KRAS4B, RHOG, ARAF, BRAF (GeneID 673), RASSF5, AFDN, PI3Kα (GeneID 5290), ELMO1 (GeneID 9844), RASSF3 (GeneID 283349), and SHOC2 (GeneID 8036). Mutants in all constructs were generated using site-directed mutagenesis. Antibodies include anti-GFP (Abcam, ab290; WB: 1:5000), anti-RAF1 (BD Biosciences, 610152; WB: 1:1000), anti-FLAG M2 (Sigma, F3165; WB: 1:1000),anti-pERK (Cell Signaling, 9101; WB: 1:1000), anti-ERK (Cell Signaling, 4695; WB: 1:1000), anti-c-MYC (Sigma, 11667149001; WB: 1:1000), anti-mCHERRY (Abcam, ab167453; WB 1:1000), anti-GAPDH (Sigma, MAB374; WB:1:1000), anti-EGFR (Cell Signaling, 2232S; WB: 1:500) HRP-conjugated anti-rabbit IgG (Cytiva, NA934; WB: 1:10000) and HRP-conjugated anti-mouse IgG (Cytiva NA931, 1:10000).

### Cell culture and rapamycin treatment

Human embryonic kidney epithelial cells (HEK 293T: ATCC CRL-3216; HEK 293A: Invitrogen) were maintained in Dulbecco’s Modified Eagle Medium (DMEM) supplemented with 10% fetal bovine serum. Cells were seeded in 6-well plates and transfected using PEI. For assay of KRAS activity, cells were starved of serum 48 hours after transfection. Rapamycin was added to media at the indicated concentrations and timepoints, after which cells were washed with PBS and lysed in 20 mM Tris-HCl (pH 7.5), 150 mM NaCl, 10% Glycerol, 1% NP-40, 5 mM MgCl_2_, 1 mM PMSF, 5 mM Okadaic acid, 1 mM Vanadate and protease inhibitor. Lysates were cleared by centrifugation and total protein concentration quantified using a BCA assay (bicinchoninic acid; Pierce). Equal amounts of lysate were loaded on SDS-PAGE gels, followed by transfer to nitrocellulose membranes for Western blotting. For this, membranes were blocked in Tris-buffered saline plus 0.1% Tween (TBST) containing 5% skim milk or 3% Bovine Serum Albumin (BSA; for phospho-specific antibodies). Following overnight incubation with primary antibodies, membranes were washed three times and blots incubated with secondary antibodies conjugated to horseradish peroxidase. Blots were treated with enhanced chemiluminescence reagent and signals were detected using a ChemiDoc system (Bio-Rad). All gel analyses were done using ImageLab (Bio-Rad). To monitor total ERK levels, anti-pERK blots were stripped using 0.7 M NaOH for 10 minutes, washed, and subsequently incubated with anti-ERK antibody.

### Generation of stable split-TEV cell lines

To derive a monocolonal cell line that stably expresses the nTEV and cTEV fragments, HEK 293A cells were first transfected with a construct expressing FLAG-FKBP-cTEV-CaaX^L190T^. Cells were grown in Zeocin (300 ng/ul) to select for cTEV expression, and these were then transduced with lentivirus produced in HEK 293T cells for expression of FLAG-FRB-nTEV. Infected cells were selected with purocycin (2 µg/ml) and serially diluted in 6-well dishes. Individual colonies were propagated and screened (by Western blot) to establish monoclonal cell lines expressing both FLAG-fused nTEV and cTEV-CaaX^L190T^ fragments at similar levels.

### Co-precipitation assays

For co-immunoprecipitation (co-IP) of protein complexes, HEK 293T cells seeded in 6-well plates were co-transfected with effector proteins (BRAF, ARAF or ELMO1) and either VENUS-tagged GTPases (KRAS or RHOG) or SPaRTa constructs (ARAF-tev-KRAS^G12V^ or Elmo2-tev- RHOG^G13D^) plus/minus rapamycin. For semi-endogenous IP with RAF1, HEK 293T cells were seeded on 10 cm plates and transfected with either VENUS alone, VENUS-KRAS^G12V^ or VENUS-ARAF-tev-KRAS^G12V^. After 48 hours, transfected cells were lysed in 20 mM Tris-HCL (pH 7.5), 150 mM NaCl, 5 mM MgCl_2_, 10% glycerol, 1% Triton X-100, 1 mM DTT, and protease inhibitor and kept on ice for 10 min. Lysates were cleared by centrifugation and incubated with pre-washed Protein G Sepharose bound with the immunoprecipitating antibody (either anti-FLAG or anti-GFP). Following 2 hours of incubation at 4°C, beads were washed three times with lysis buffer, reconstituted in sample buffer, separated by SDS-PAGE and transferred to nitrocellulose membranes for Western blot analysis.

### Microscopy

Cells were split on ethanol sterilized coverslips in 6 well dishes and incubated at 37°C in 5% CO2 for 24 hours. Following transient expression of constructs for 48 hours (and overnight starvation for KRAS activity assays), cells were washed with PBS and fixed with 4% Paraformaldehyde (VWR). Coverslips were washed with PBS and incubated with Hoechst (1:2000) for nucleic acid staining. Following a final wash with PBS and ethanol, coverslips were mounted on slides with the Prolong Diamond antifade mountant (Life Technologies) and dried for 24 hours before acquisition. Imaging of cells was performed using a laser scanning LSM-880 microscope (Zeiss). All images were taken with a 63x objective. 12 to 15 z-stacks were acquired (0.25 um thickness) for each image and were merged by an XY orthogonal projection with the Zen lite 2.3 software (Zeiss).

### Fractionation

HEK 293A cells stably expressing FLAG-FRB-nTEV and FLAG-FKBP-cTEV-CaaX^L190T^ were transfected to transiently express CHERRY-tagged effectors with VENUS alone, VENUS-tagged GTPases, or VENUS-SPaRTa constructs. 24 hours post-transfection, cells were washed once with PBS and serum starved for 18 hours. SPaRTa samples were treated for the specified timepoints with 50 nM rapamycin. As required, KRAS^WT^ expressing cells were treated with EGF (100 ng/ml) for 5 minutes. Following treatment, cells were washed with PBS, collected with a scraper and gently pelleted by centrifugation. PBS was carefully removed, and the cell pellet resuspended in 500 µl of ice-cold homogenization buffer (50 mM HEPES-NaOH (pH 7.0), 0.25 M sucrose, 78 mM KCl, 4 mM MgCl_2_, 8.4 mM CaCl_2_, 10 mM EGTA, 1 mM DTT, 1 mM PMSF and protease inhibitor). Lysates were subsequently passed through a 27-gage needle 30X. Cell death was verified by loading a mixture of 10 µl lysate and 10 µl Trypan Blue in a cell counting chamber and assessing lysis with a Countess Automated Cell Counter. Homogenates were cleared by centrifugation at 1,500xg for 10 minutes at 4°C. Supernatants were transferred carefully without disturbing the pellet to a chilled Beckman tube (#343776) and fractionated by ultracentrifugation (Sorvall) at 100,000xg for 45 minutes. Supernatants (cytosol fraction) were transferred to a fresh Eppendorf tube without disturbing the membrane fraction. Remaining pellets were washed with homogenization buffer by gently pipetting the wall of the tube. All traces of buffer were removed, and membrane pellets were resuspended in 100 µl of buffer. For crosslinking, 1 mM DSS (Sigma S1885) was added to cells following starvation, treatment and resuspension in PBS. Samples were incubated on a nutator at room temperature for 15 minutes. Crosslinker was quenched using 20 mM Tris-HCl (pH 7.0) for 10 minutes at room temperature, and cells centrifuged at 1,200 rpm. All traces of PBS/DSS/Tris were removed, and the cell pellet resuspended in 500 µl of homogenization buffer for subsequent fractionation.

## Acknowledgements

This work was supported by grants from the Canadian Institutes for Health Research (CIHR, to M.J.S.) and the National Science and Engineering Council of Canada (NSERC, to M.J.S.). M.J.S. holds a Canada Research Chair (CRC) in Cancer Signalling and Structural Biology. S.S. was supported by scholarships from the Fonds de recherche du Québec (FRQ).

## Data Availability statement

All data is included in the manuscript. Requests for materials should be sent to Matthew Smith.

---

**Supplementary Figure S1.**
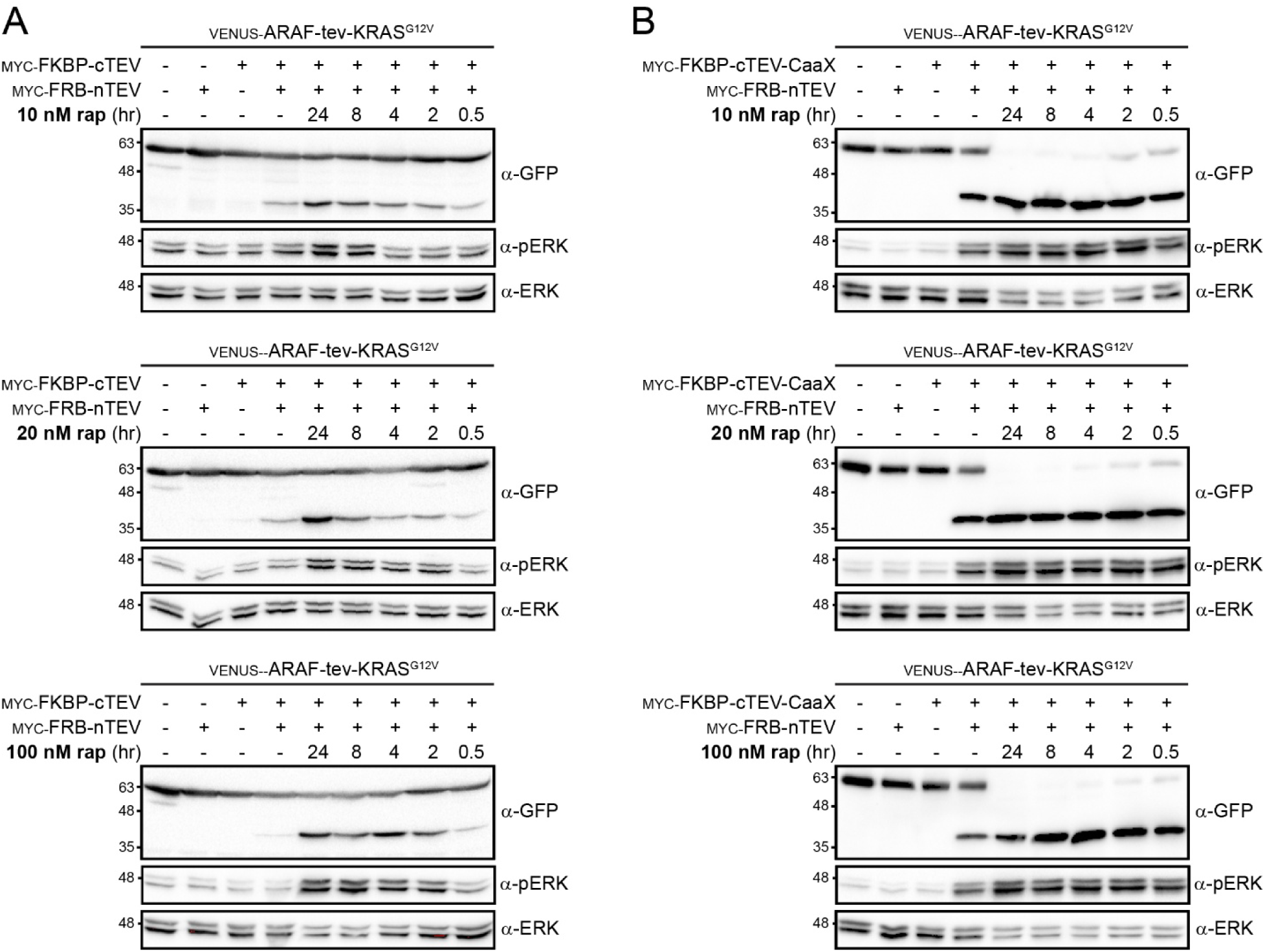
Influence of rapamycin concentration on cleavage efficiency. (A) Timecourse of occluded VENUS-ARAF-tev-KRAS^G12V^ activation induced by rap over 24 hours. The occluded GTPase was co-expressed with nTEV and cTEV and stimulated with either 10 nM (top), 20 nM (middle), or 100 nM (bottom) rap for the indicated timepoints. Blotting for pERK reveals downstream signalling from KRAS^G12V^, evident once the tethered RBD is released by TEV-linker cleavage. Production of VENUS-ARAF RBD reports on TEV activity. (B) Activation of occluded VENUS-ARAF-tev-KRAS^G12V^ when co-expressed with nTEV and cTEV-CaaX is independent of rap concentration. Cells were stimulated with 10 nM (top), 20 nM (middle), or 100 nM (bottom) rap for the indicated timepoints. The production of VENUS-ARAF RBD and blotting for pERK/ERK report on activity, which becomes significantly decoupled from rap in this system.

**Supplementary Figure S2.**
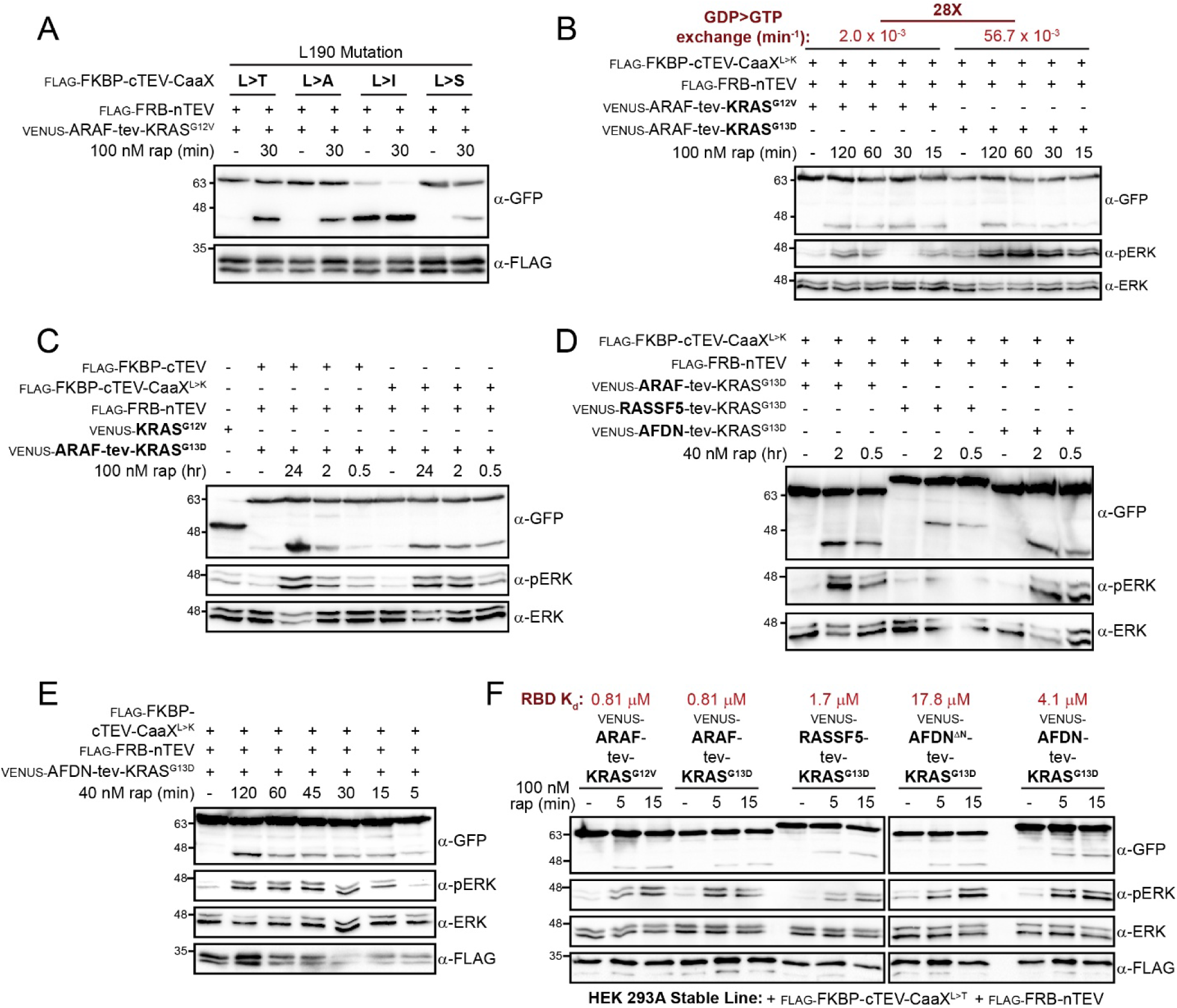
Multiple approaches to optimize the occluded KRAS construct. (A) Restoring cTEV-CaaX responsiveness by introducing mutations at the TEV reconstitution interface. FKBP-cTEV mutants L190T, L190A, L190I and L190S were co-expressed with FRB-nTEV and occluded VENUS-ARAF-tev-KRAS^G12V^. Cells were stimulated with 100 nM rap for 30 min and cleavage efficiency monitored by observing production of VENUS-ARAF RBD. (B) Comparison of MAPK stimulation from occluded KRAS constructs with activating mutation G12V versus G13D. The intrinsic exchange activity (GDP to GTP) of these mutants differs by 28-fold, with the G13D mutant exhibiting more rapid exchange (labelled red, top). ARAF RBD occluded constructs were co-expressed with cTEV-CaaX^L190K^ and nTEV. 100 nM of rap was added for the indicated timepoints. Production of VENUS-ARAF RBD verifies cleavage, and pERK/ERK blots reveal efficiency of each KRAS mutant at stimulating MAPK activity. (C) The KRAS mutant G13D maintains RBD occlusion and rapid pERK induction following rap addition, with levels of MAPK activation higher than observed for prolonged expression of VENUS-KRAS^G12V^ alone. The VENUS-ARAF-tev-KRAS^G13D^ construct was co-expressed with nTEV and either the archetype cTEV fragment or cTEV-CaaX^L190K^. Cleavage efficiency following addition of 100 nM rap for the indicated timepoints is assayed both by the production of VENUS- ARAF RBD and induction of pERK. (D) Tethering of RBDs with diverse binding affinities for KRAS to optimize RBD diffusion following TEV-linker cleavage. RBD domains from ARAF (K_d_=0.81 µM), RASSF5 (1.7 µM) or AFDN (4.1 µM) were tethered to KRAS^G13D^. Constructs were co-expressed with nTEV/cTEV-CaaX^L190K^ and 40 nM rap added for the indicated timepoints. VENUS-tagged RBDs and pERK induction report on cleavage efficiency. (E) Full cleavage timecourse of VENUS-AFDN-tev-KRAS^G13D^ co-expressed with nTEV and cTEV-CaaX^L190K^ (rap addition over 5-120 min). pERK/ERK immunoblots report on KRAS signalling activity, reflecting release of the occluding RBD domain. Expression of TEV fragments is confirmed by anti-FLAG blot (bottom). (F) Comparison of occluded construct activation in HEK 293A cells stably expressing FKBP-cTEV-CaaX^L190T^ and FRB-nTEV. Either KRAS^G12V^ (occluded by ARAF RBD) or KRAS^G13D^ (occluded by RBDs from ARAF, RASSF5 or AFDN) were expressed and TEV reconstituted by addition of 100 nM rap for 5 or 15 min. The previously measured affinities (K_d_) of each RBD for KRAS are listed at top (red). A shortened AFDN domain with very low affinity (17.8 µM) was included. Cleavage is confirmed by production of VENUS-RBDs and signalling efficiency by induction of pERK. A FLAG blot verifies expression of the TEV fragments.

**Supplementary Figure S3.**
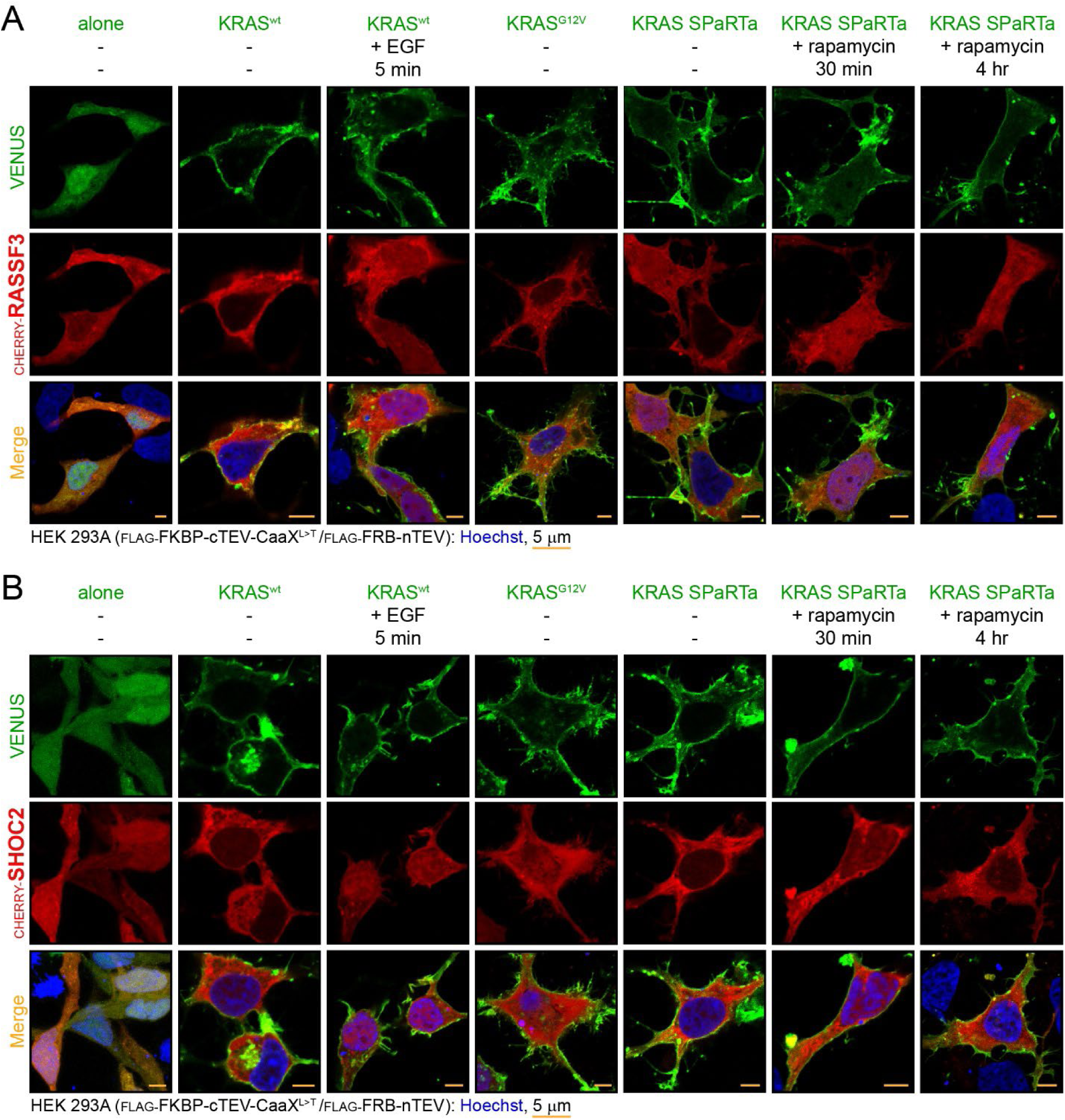
The localization of RASSF3 and SHOC2 in the cytosol is unaffected by KRAS activation. (A) CHERRY-RASSF3 is localized in the cytosol of starved HEK 293A cells expressing VENUS alone. EGF stimulation (50 ng/ml for 5 minutes in cells expressing VENUS-KRAS^WT^), constitutively active VENUS-KRAS^G12V^, or rap-induced activation of KRAS SPaRTa all fail to recruit RASSF3 to the PM. Scale bar is 5 µm; Hoeschst-stained nuclei are blue. (B) The distribution of CHERRY-SHOC2 in the cytosol of HEK 293A cells is unchanged by KRAS activation. Addition of growth factor (50 ng/ml EGF for 5 minutes in cells expressing VENUS-KRAS^WT^), co-expression of SHOC2 with constitutively active VENUS-KRAS^G12V^, or activation of KRAS SPaRTa for the indicated timepoints (with 40 mM rap) had no impact on SHOC2 localization. Scale bar is 5 µm; Hoeschst-stained nuclei are blue.

**Supplementary Figure S4.**
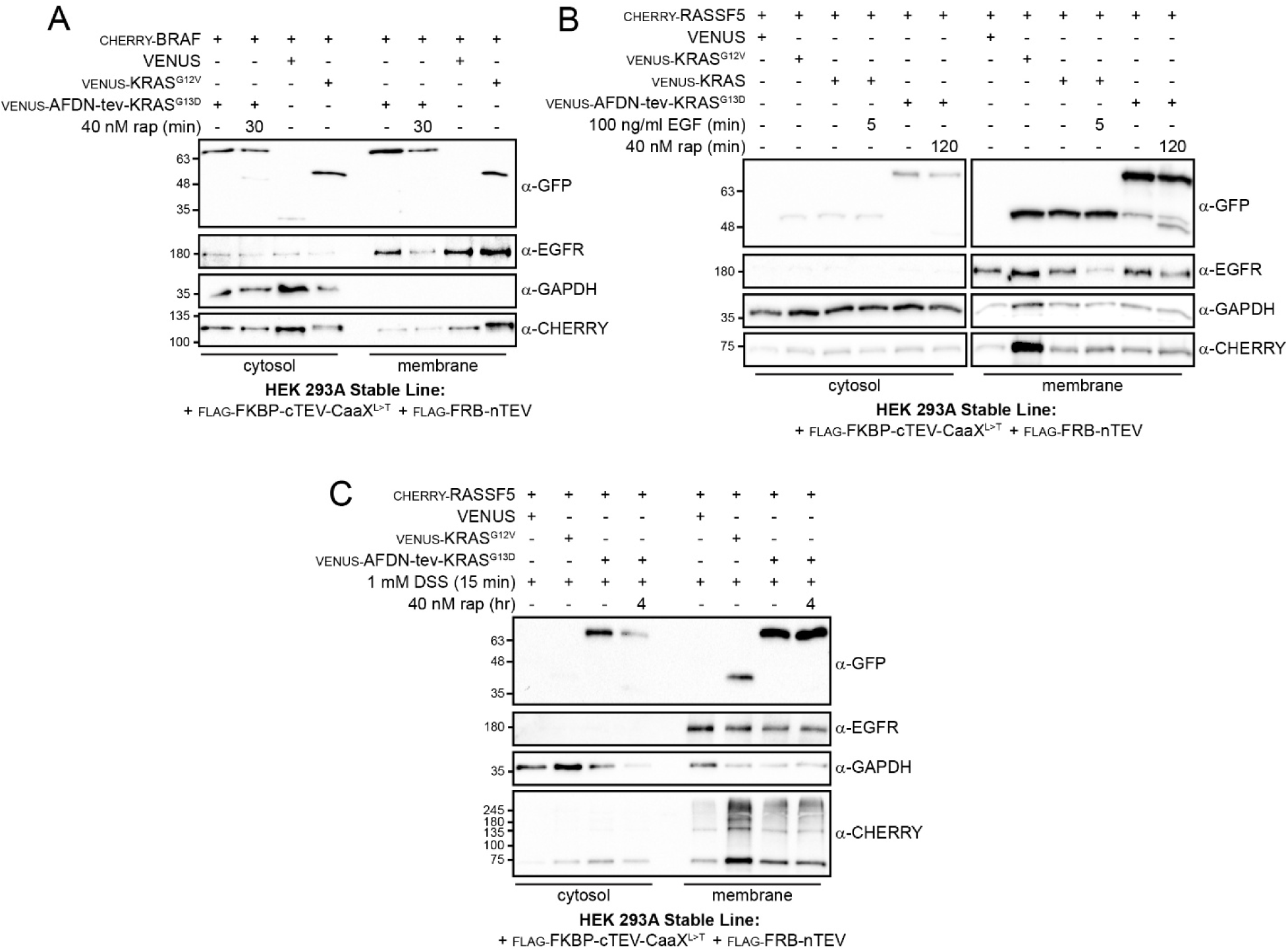
Transient recruitment of effectors to the PM by KRAS activation is not detectable using fractionation. (A) Comparison of cytosol and membrane fractions from cells expressing CHERRY-BRAF and either a VENUS alone control, activated VENUS-KRAS^G12V^ or the VENUS-AFDN-tev-KRAS^G13D^ SPaRTa construct. Occluded KRAS was activated by addition of 40 nM rap for 30 min. Only a 48-hr expression of the constitutively on KRAS^G12V^ enriched BRAF in the membrane fraction. GAPDH (predominantly cytosolic) and EGFR (predominantly PM-localized) are controls. (B) CHERRY-RASSF5 is enriched in the membrane fraction only by extended expression of VENUS-KRAS^G12V^. Neither stimulation of cells expressing VENUS-KRAS^WT^ (with 100 ng/ml EGF; 5min) or activation of the KRAS SPaRTa (with 40 nM rap; 120 min) showed increased RASSF5 in membrane fractions. Cleavage is verified by the presence of VENUS-AFDN RBD. GAPDH (predominantly cytosolic) and EGFR (predominantly PM-localized) are fractionation controls. (C) Crosslinking with DSS (1 mM for 15 min) following activation of KRAS SPaRTa does not change RASSF5 distribution in the cytosolic/membrane fractions. VENUS-AFDN-tev-KRAS^G13D^ was activated by addition of 40 nM rap for 4 hours. Significant laddering of CHERRY-RASSF5 is visible on Western blots upon incubation with DSS, but there is no enrichment in the membrane fraction produced by SPaRTa activation over controls. As expected, constitutively active VENUS-KRAS^G12V^ did increase RASSF5 levels in the membrane fraction.

**Supplementary Figure S5.**
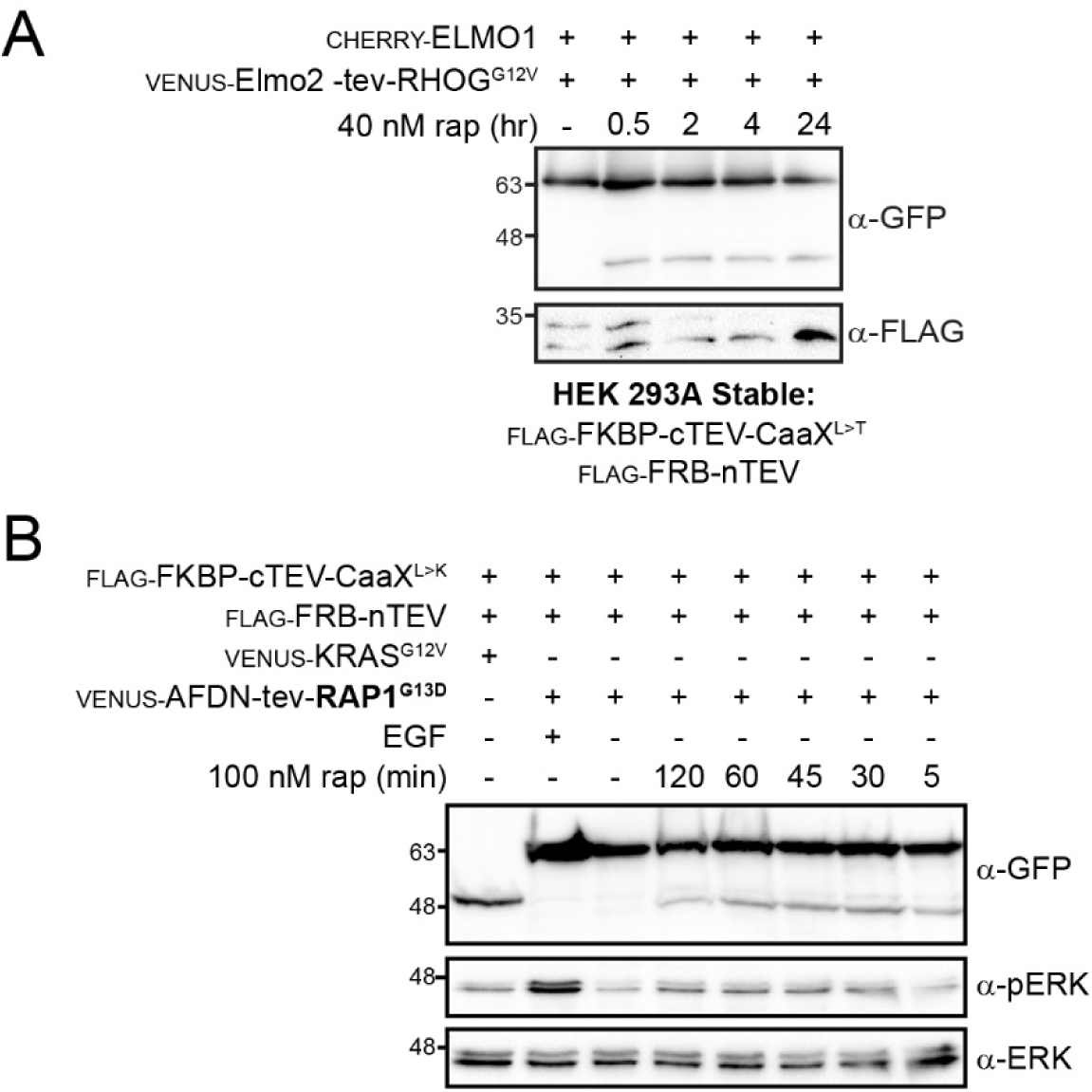
SPaRTa approach for diverse GTPases in the RAS superfamily. (A) Verification of rap-induced RHOG SPaRTa cleavage for confocal microscopy experiments. Following removal of coverslips from culture dishes, remaining cells were lysed and assayed for cleavage. Experiment included addition of 40 nM rap for the indicated timepoints. Presence of VENUS-Elmo2 RBD verifies activation, and an anti-FLAG blot shows TEV fragments. (B) Timecourse of MAPK activation downstream of a RAP1 SPaRTa (VENUS-AFDN-tev-RAP1^G13D^). The occluded construct was co-expressed with nTEV and cTEV-CaaX^L190K^. Cleavage was induced by addition of 100 nM rap for the indicated timepoints. The production of VENUS-AFDN RBD verified cleavage, with MAPK induction assayed by pERK stimulation.

